# Deep-learning enables proteome-scale identification of phase-separated protein candidates from immunofluorescence images

**DOI:** 10.1101/636738

**Authors:** Chunyu Yu, Boyan Shen, Qi Huang, Minglei Shi, Kaiqiang You, Congying Wu, Yang Chen, Tingting Li

**Affiliations:** Department of Biomedical Informatics, School of Basic Medical Sciences, Peking University Health Science Center, Beijing 100191, China; Institute of Systems Biomedicine, School of Basic Medical Sciences, Peking University Health Science Center, Beijing 100191, China; MOE Key Laboratory of Bioinformatics; Bioinformatics Division and Center for Synthetic & Systems Biology, TNLIST; School of Medicine, Tsinghua University, Beijing 100084, China

## Abstract

Intrinsically disordered region (IDR) analysis has been widely used in the screening of phase-separated proteins. However, the precise sequences determining phase separation remain unclear. Furthermore, a large number of phase-separated proteins that exhibit relatively low IDR content remain uncharacterized. Phase-separated proteins appear as spherical droplet structures in immunofluorescence (IF) images, which renders them distinguishable from non-phase-separated proteins. Here, we transformed the problem of phase-separated protein recognition into a binary classification problem of image recognition. In addition, we established a method named IDeepPhase to identify IF images with spherical droplet structures based on convolutional neural networks. Using IDeepPhase on proteome-scale IF images from the Human Protein Atlas database, we generated a comprehensive list of phase-separated candidates which displayed spherical droplet structures in IF images, allowing nomination of proteins, antibodies and cell lines for subsequent phase separation study.

## Introduction

How cells segregate cellular components in an appropriate temporal and spatial manner remains a fundamental question in cell biology. In addition to classic membrane-bound organelles, membraneless organelles offer a flexible way to regulate location and concentration of these cellular components. Under specific physiological conditions, multivalent molecules such as proteins and nucleic acids, termed scaffolds, undergo polymerization and recruit a class of proteins and nucleic acids, termed clients, and assembles into small compartments. Such compartmentalization allows components to be exchanged with a cell’s environment in a tightly controlled manner ^1^. These compartments are also called membraneless organelles, biomolecular condensates or phase-separated compartments ^1^. Common membraneless organelles include nucleoli, PML-body, stress granules and P body. It was recently shown that liquid–liquid phase separation (LLPS) underlies the formation of membraneless organelles ^2, 3^. As a consequence, LLPS is now viewed as determinant of various pathological processes underlying, for instance, neurodegenerative diseases ^1, 2, 4^ as well as oncogenesis. A recent report showed that the prion-like domain of EWS-FLI1 fusion protein phase separates and recruits BAF chromatin-remodeling complexes to tumor-specific enhancers, thereby activating the transcriptional events of Ewing’s sarcoma ^5^.

Understanding the principles underlying the formation of membraneless organelles is crucial for investigating the physiology and pathophysiology of various biological processes. Furthermore, identifying the proteins linked to LLPS represents a critical first step towards characterizing membraneless organelles. Several recent studies found that many proteins involved in LLPS contain a high proportion of intrinsically disordered regions (IDRs); furthermore, these regions usually encompass low-complexity sequence regions (LCRs) ^1,4,6^. Therefore, IDR content analysis is often utilized in bioinformatics screening of phase-separated proteins ^7^. With programs predicting disorders such as D2P2 ^8^ and MobiDB ^9^, proteins with high proportion of IDRs be screened out. In addition to IDR length and number, sequence patterns like prion-like LCRs are also related to phase separation. Therefore, several phase-separation predictors further predict whether specific sequence patterns were included in the screened proteins, e.g. Pi-Pi predictor for pi-pi contacts prediction ^10^, PLAAC for prion-like domain prediction ^11^ and ZipperDB for fibril-forming segment prediction ^12^. While recent studies suggest that the length, number, and sequence pattern of IDRs are involved in fine-tuning their phase behavior, our understanding of how IDRs determine phase separation remains limited ^13^. Furthermore, IDR content analysis tools are not suitable for phase-separated proteins with relatively low IDR content, including SOS1 in T cell microclusters ^14^, ELF2 in stress granule^15^, mTOR in PML body ^16^ and PGL granule of *C.elegans* ^17^.

One reliable criterion for phase separation is when spherical structures are formed and fuse with one another ^13^. The most common and convenient approach to identify these spherical droplet structures through IF images. With this method, marker proteins are fluorescently labeled to trace certain phase-separated compartment formed *in vitro* as well as in cells. A recent study analyzed clustered IF images of normal and perturbed cells to characterize the regulatory function of 1354 human genes on the size, number and morphology of typical membraneless compartments ^18^. The specific droplet structure in the IF images is a typical feature which distinguish phase-separated proteins from other proteins ^13,19^. In another recent study, deep learning models based on IF images have been successfully established for the identification of protein subcellular locations ^20^. Here, we set out to investigate whether deep learning based on IF images is an efficient way to discriminate proteins involved in phase separation.

## Results

### Overview of IDeepPhase method

We collected 96 experimentally confirmed human phase-separated proteins by retrieving from the literature (Supplementary Table 1). IF images of these proteins were downloaded from the Cell Atlas of Human Protein Atlas database (https://www.proteinatlas.org/humanproteome/cell) ^21^. Proteins involved in phase separation compartments usually displayed spherical droplet structures in IF images ^2, 13, 19^. Consequently, identification of the proteins to be involved in phase separation was transformed into a binary classification problem of image recognition, whose goal was to identify IF images with spherical droplet structures. Since the formation of phase separation depended on different conditions ^1, 2
^, we manually picked up IF images which display droplets with higher fluorescence intensity than background as positive samples (Supplementary Table 2). Negative samples were randomly selected from all downloaded IF images, after filtering samples corresponding to the collected phase-separated proteins (Supplementary Table 2).

Using positive and negative samples, we next constructed a classifier called IDeepPhase based on a convolutional neural network (CNN). As shown in Fig. 1a, a two-step training approach was used to build IDeepPhase. In the first step, the positive and negative sample sets were separated into training and test sets. The positive training set includes 195 IF images and the negative training set includes an equal-sized IF images (Supplementary Table 2). The remaining 79 IF images and an equal-sized negative set were used as test dataset (Supplementary Table 2). To augment the sample size of the training and test datasets, each IF image was cropped into 4 non-overlapping pieces, which increased the number of samples 4-fold. The CNN model of the first step was trained with the enlarged positive and negative training sets. In the second step, we used the CNN classifier trained in the first step to score all IF images downloaded from Human Protein Atlas database. Each IF image was cropped into 4 non-overlapping pieces and finally 334692 cropped images were scored. The cropped images scoring in the top 1000 were manually labeled and added to the positive and negative training sets. Next, an extended training set with include 1241 positive samples and 1253 negative samples were built. The CNN network was then retrained with this extended training set, and the test dataset was the same as the first stage. The final AUC of IDeepPhase on the test set was 0.91 (Fig.1b), indicating that IF images with droplet structures can be discriminated effectively with our deep learning approach.

**Fig. 1.**
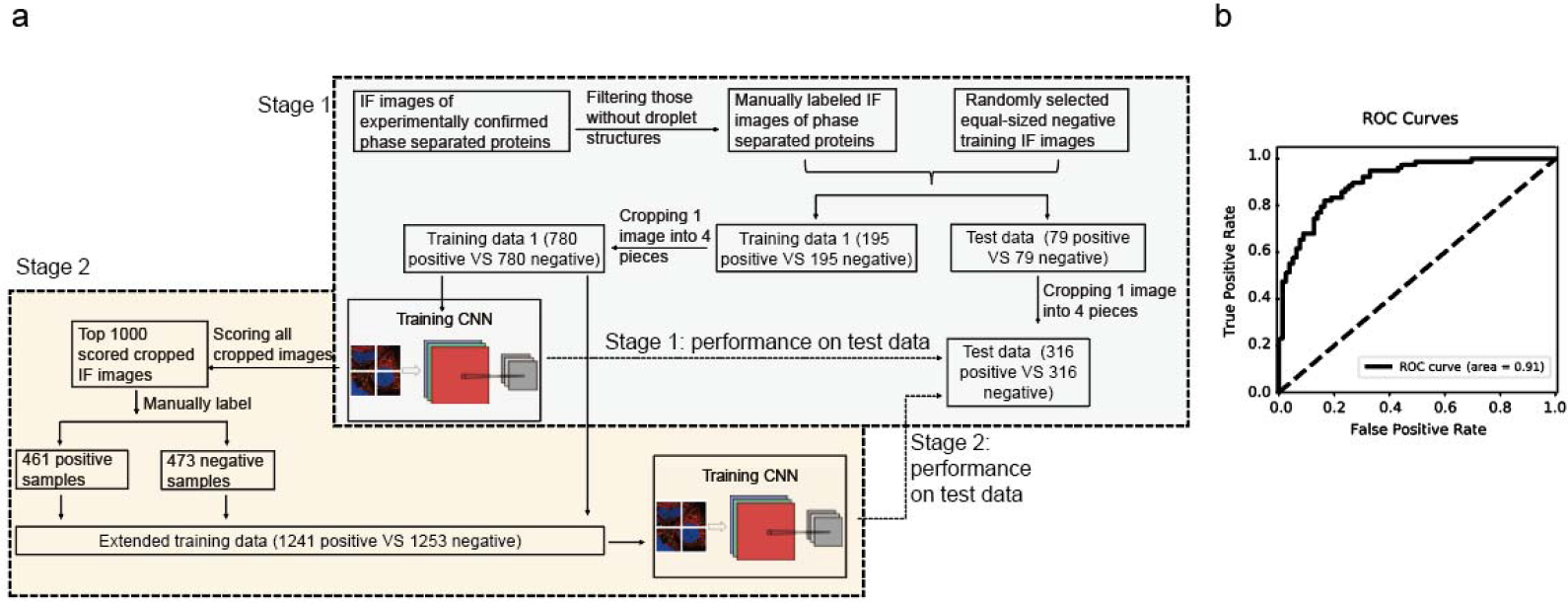
The IDeepPhase frame work and performance. **a**. A deep learning approach for identifying droplet structures on IF images. **b**. Receiver operating characteristic (ROC) curve of classification on the test set.

### Identification of IF images with droplet structures

Here we built a CNN classifier called IDeepPhase, which discriminates IF images that display droplet structures. Using IDeepPhase, 334692 cropped IF images were scored. For each protein, multiple scores were generated since at least four cropped images were available. The highest score was taken as the final score of the protein (Supplementary Table 3). The distribution of IDeepPhase scores (IDP scores) of all the proteins are shown in Fig. 2a.

**Fig. 2.**
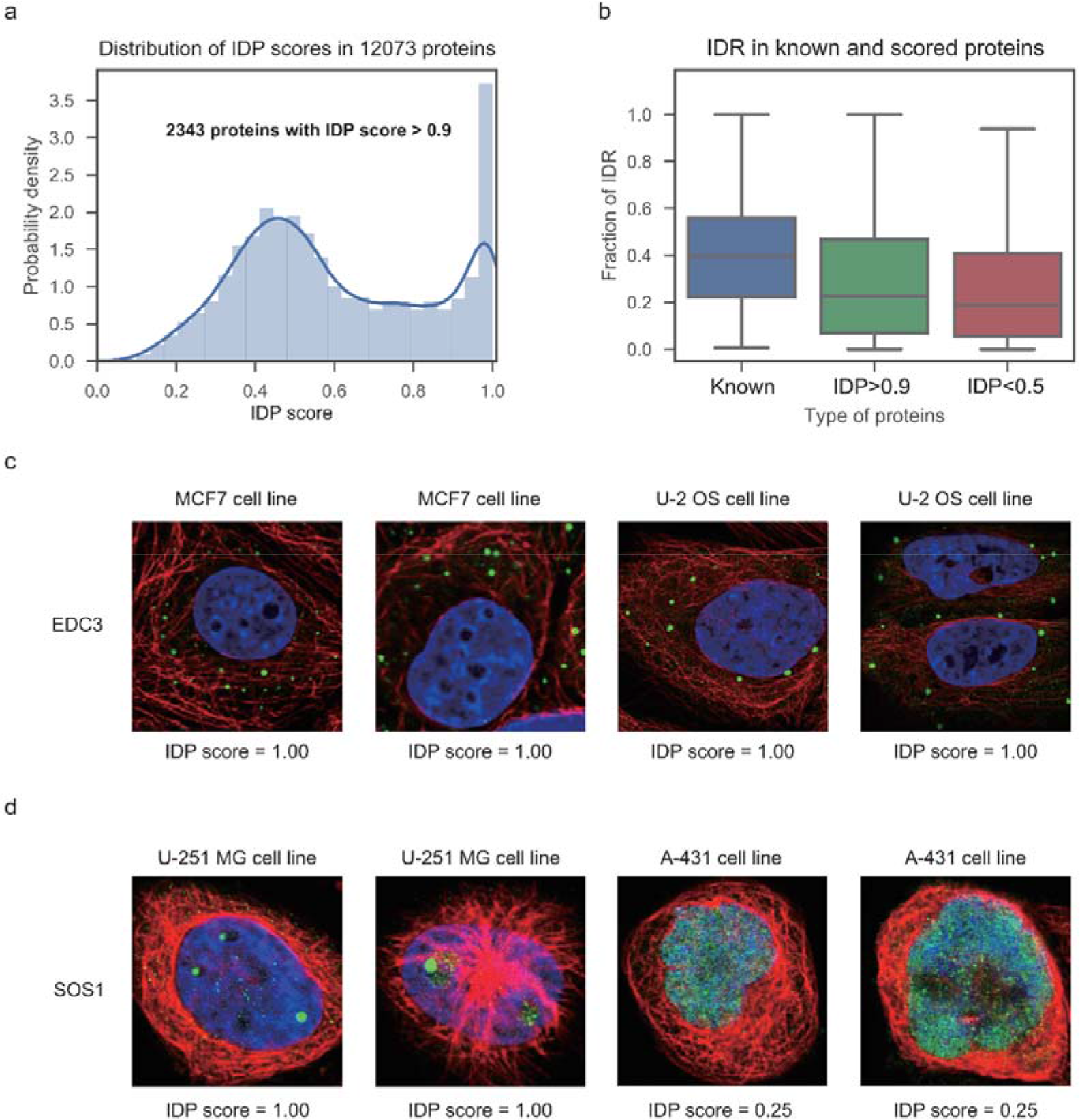
Systematic identification of IF images with droplet structures. **a**. Distribution of IDP scores for 12073 proteins. **b**. Fraction of intrinsically disordered regions (IDRs) in known phase-separated proteins, proteins with IDP scores higher than 0.9 (IDP > 0.9) and less than 0.5 (IDP < 0.5). **c**. IF images of EDC3 in MCF7 cell line and U-2 OS cell line. **d**. IF images of SOS1 in U-251 MG cell line and A-431 cell line.

Previous studies found that proteins involved in phase separation usually possess a high proportion of IDRs ^1, 4, 6^. In agreement with this phenomenon, the IDR content of proteins with IDP scores higher than 0.90 was significantly higher than proteins with IDP scores less than 0.5 (One-tailed Mann-Whitney U test, p = 7E-8, Fig. 2b). The difference between the highly scored proteins and the known phase-separated proteins was significant as well (One-tailed Mann-Whitney U test, p = 5E-7, Fig. 2b). One possible explanation is that researchers who searched phase-separated candidates for further study prefer proteins containing a higher proportion of IDRs. It is well established that a number of phase-separated proteins exhibits relatively low IDR content. For example, the EDC3 protein has been shown to locate in P-bodies ^22,23^. According to the prediction of D2P2, about 20% of EDC3 peptide sequence was consisted of IDRs. As shown in Fig. 2c, the IDP score of EDC was 1 in both MCF7 and U-2 OS cell lines. Another example was SOS1, which has been demonstrated to form phase separation in the process of T cell receptor signal transduction ^14^. According to the prediction of D2P2, about 25% of SOS1 peptide sequence was consisted of IDRs. As shown in Fig. 2d, the IDP score of SOS1 was 1 in U-251 MG cell line. Together, these results suggested that proteins with relative low IDR content can be discriminated by IDeepPhase.

It should be noted that IDeepPhase was built to discriminate IF images which displayed droplet structures. In addition, the formation of droplet structures is dependent on the concentration, solubility, valency and post-translational modification of the protein ^1^. Therefore, phase-separated proteins displaying diffuse signals cannot be identified by IDeepPhase. As shown in Fig. 2d, SOS1 displayed a droplet structure in U-251 MG cell line, whereas the signal was more diffused in A-431 cell line. Another example was EWSR1, whose IF images were generated in three cell lines A-431, U-2 OS and U-251MG. However, the droplet structure of EWSR1 was only detected in U-2 OS cells (Supplementary Fig. 2). Therefore, the antibodies and the cell lines which generated the highest scores are provided in Supplementary Table 3.

### Components of both intranuclear and extranuclear membraneless organelles can be discriminated by IDeepPhase

To further investigate whether the IDP scores extracted from the GFP-tagged IF images of specific protein indicate phase separation potential, we compared the IDP scores with the membraneless organelle annotations from the Swiss-Prot database and the components of membraneless organelles generated by high-throughout approaches. In these comparisons, proteins used as training samples were removed (Supplementary Table 4).

In nucleoli, multiple liquid phase subcompartments may execute different processing steps in ribosome biogenesis, and these subcompartments serves similar to different workshops on an assembly line ^24^. The nucleolar proteome includes hundreds of proteins, which are considered to segregate into at least three distinct compartments where three main steps of ribosome biogenesis mainly take place: the fibrillar center of rDNA transcription, the dense fibrillar component of rRNA processing and the granular component of ribosome assembly. Proteins POLR1E, FIB1 and NPM1 have been used as markers of the above three liquid phase subcompartments. The IDP scores of POLR1E, FIB1 and NPM1 are all 0.99, suggesting that IDeepPhase identified known phase-separated proteins (Supplementary Table 4). To further investigate whether most of the nucleolar proteins can be discriminated by IDeepPhase, proteins annotated to locate in nucleoli were extracted from Swiss-Prot database (Supplementary Table 4). As shown in Fig. 3a, IDP scores of nucleoli proteins were significantly higher than other proteins excluding nucleoli components (One-tailed Mann-Whitney U test, p = 7E-24, Fig. 3a).

**Fig. 3.**
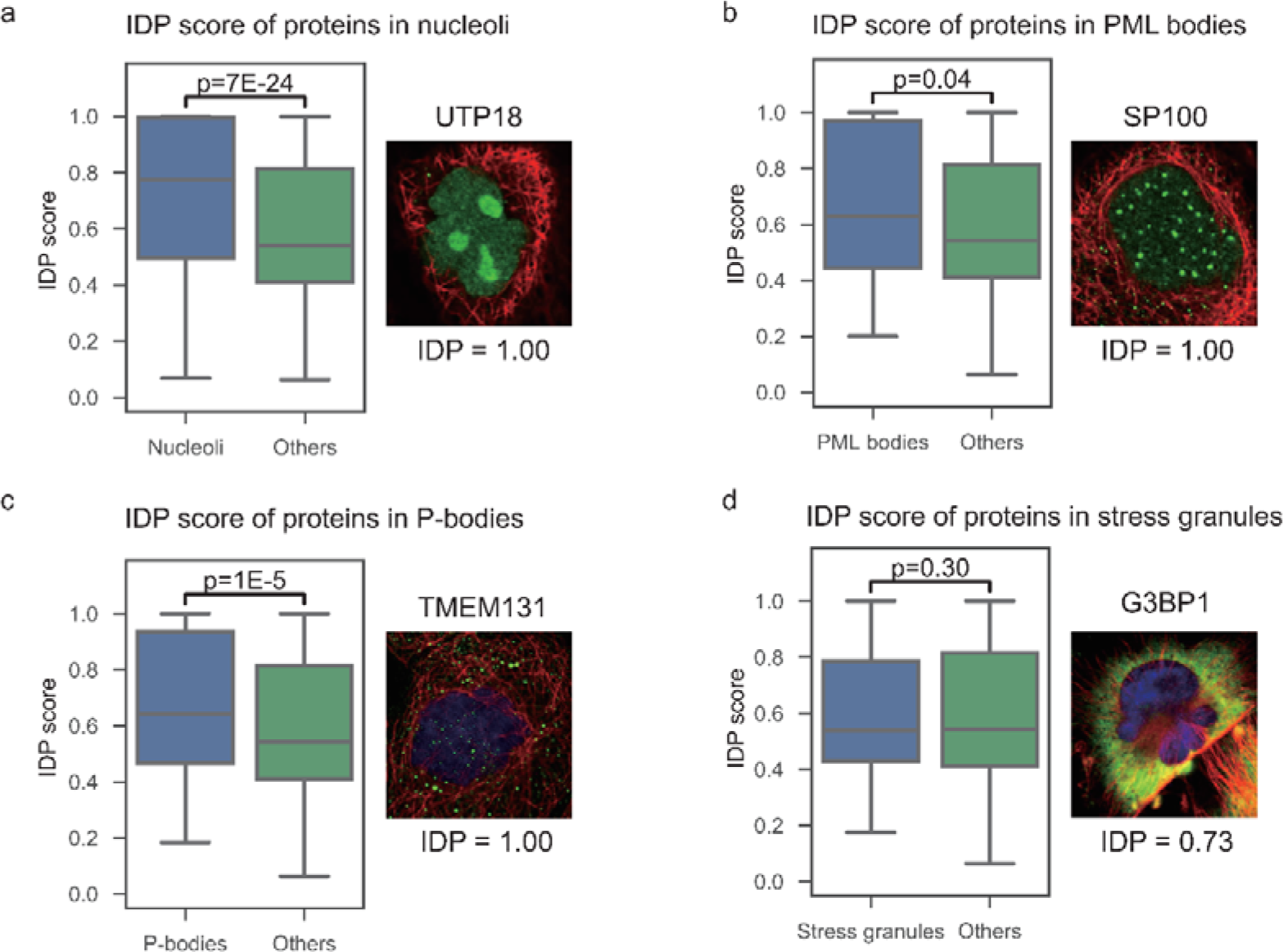
Comparisons of IDP scores for proteins localized within and without certain phase-separated component. nucleoli (**a**), PML bodies (**b**), P-bodies (**c**), and stress granules (**d**).

The PML body is another typical phase-separated nucleolar subcompartment. Proteins ATRX, MRE11, HIPK3, RPAIN and TDP2 were previously demonstrated to be the components of PML body ^25^. As shown in Supplementary Fig. 2, their IDP scores are identified as 0.91, 0.98, 0.91, 0.95 and 0.99 respectively. IDP scores of PML body components extracted from Swiss-Prot database were significantly higher than other proteins excluding PML-body components (One-tailed Mann-Whitney U test, p = 7E-24, Fig. 3b; Supplementary Table 4). Furthermore, as shown in Fig. 3a and 3b, both the components of nucleoli (UTP18) and PML body (SP100) displayed droplet structures. The size of nucleoli droplets was significantly larger than PML body droplets, and nucleoli droplets are also less likely to be standardly spherical than PML body droplets.

In addition to these two intranuclear membraneless organelles, we further investigated whether components of extranuclear membraneless organelles could be discriminated by IDeepPhase. A systematic *in vivo* analysis of proximity-dependent biotinylation (BioID) was recently conducted to identify P body components ^26^. A second P-body proteome was analyzed using a fluorescence-activated particle sorting (FAPS) method, to find proteins that are located in P-body ^23^. After filtering all proteins which were used as training samples as well as all proteins without IDP scores, we obtained 140 proteins (Supplementary Table 4). As shown in Fig. 3c, IDP scores of 140 P-body components were found to be significantly higher than other proteins excluding P-body components (One-tailed Mann-Whitney U test, p=1E-5, Fig. 3c).

Components of stress granules had been identified by proteomic analysis as well ^27^. After filtering all proteins which were used as training samples as well as all proteins without IDP scores, we obtained 257 proteins (Supplementary Table 4). However, IDP scores for 257 stress granule components were not significantly higher than other proteins excluding stress granule components (Mann-Whitney U test p-value 0.49, Fig. 3d). Stress granules are phase-separated organelles which appear to protect RNAs when cells are stressed ^28^. One recent report showed that without exogenous stress, the stress granule markers G3BP1 occur as diffuse signals throughout the cytoplasm ^26^. In agreement with this study, G3BP1 also displayed diffuse cytosolic signals in the downloaded IF images from the Human Protein Atlas Database (Fig. 3d), because components of stress granules cannot be identified from IF images generated without exogenous stress.

Taken together, these analyses clearly demonstrated that IDeepPhase was highly effective in extracting putative phase-separated components of both intranuclear and extranuclear membraneless organelles.

### Intranuclear proteins with high IDP scores possess typical features of phase-separated components

To further investigate whether proteins with high IDP scores possess typical features of phase-separated proteins, we selected 2343 proteins which received IDP scores higher than 0.9 for sequential and functional preferences analysis.

Unlike in the nucleus, extranuclear membrane-bounded organelles like endosomes or lysosomes display spherical structures similar to membraneless organelles. To investigate the influence of extranuclear membrane-bounded organelles on the prediction results of IDeepPhase, we separated 2343 proteins into two groups according to the location annotations of the Human Protein Atlas database: 1467 proteins whose locations include the nucleoli, nucleoli fibrillar center, nuclear bodies, nuclear speckles, nucleoplasm or nucleus; 876 proteins which were only annotated to be located in the extranuclear organelles.

Of 1467 intranuclear proteins, we identified 16% (235/1467) with IDR proportions higher than 0.5; while of 876 extranuclear proteins, we only identified 5% (44/876) with IDR proportions higher than 0.5 (Fig. 4a). In addition to the enrichment of IDRs, another sequential feature is that low complexity regions (LCRs) are enriched in phase-separated proteins. To evaluate the enrichment of LCRs, we downloaded the LCRs of 2343 proteins from the LCR-eXXXplorer database ^29^. For 1467 intranuclear proteins, the LCR contents of proteins with IDP scores higher than 0.90 were significantly higher than for proteins with IDP scores lower than 0.5 (One-tailed Mann-Whitney U test, p = 0.0009, Fig. 4b). In agreement with our LCR content analysis, the domain analysis shown in Fig. 4c indicate that compositionally biased regions are also enriched in 1467 intranuclear proteins. In comparison, for 867 extranuclear proteins, the LCR contents of proteins with IDP scores higher than 0.90 were not significantly higher than proteins with IDP scores lower than 0.5 (One-tailed Mann-Whitney U test, p = 0.12, fig. 4b).

**Fig. 4.**
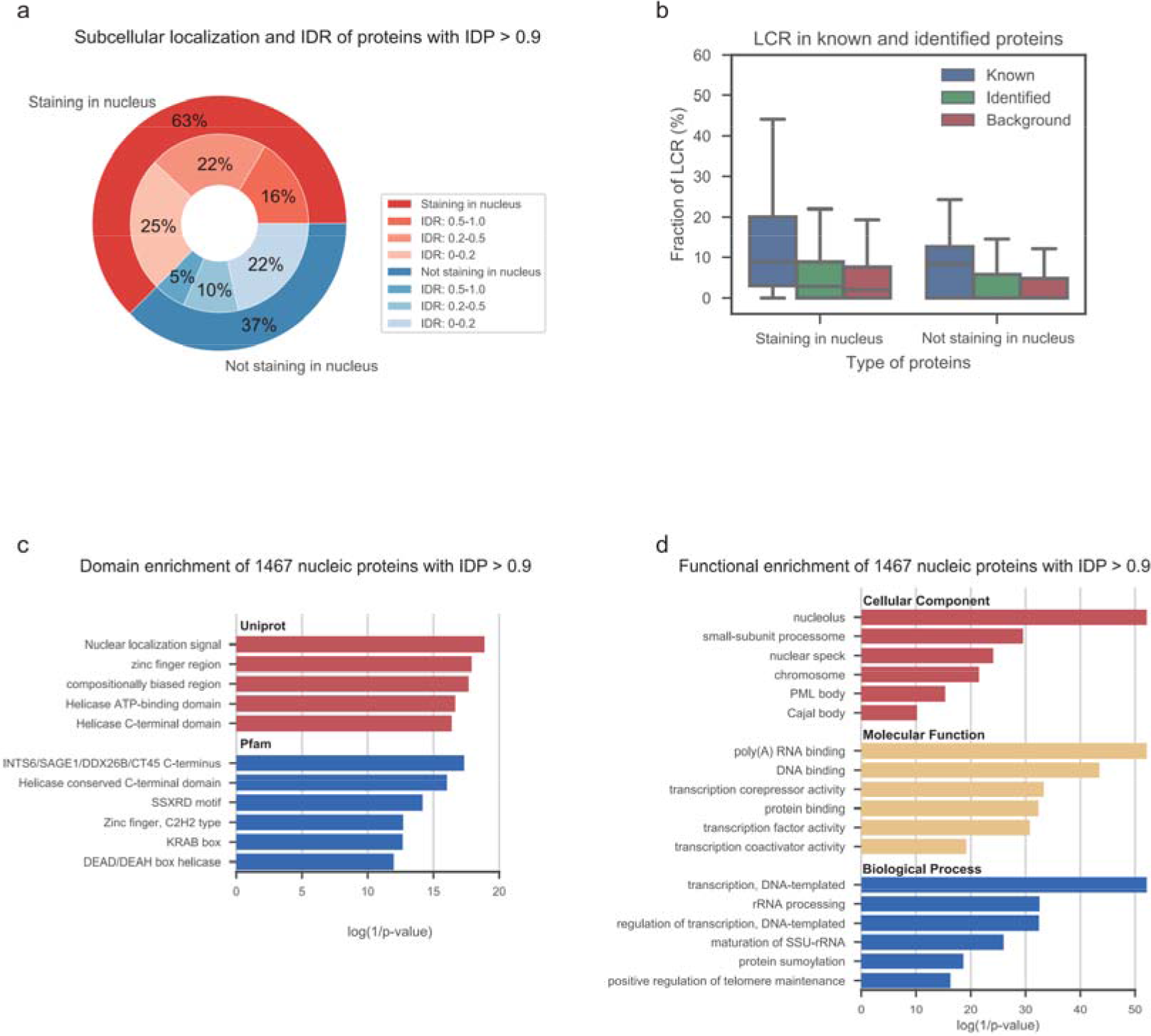
Sequential and functional preference of proteins with IDP scores higher than 0.9. **a**. IDR content summary of proteins with IDP scores higher than 0.9 staining in and out nucleus. **b**. Fraction of low-complexity regions (LCRs) in known phase-separated proteins, intranuclear proteins with IDP scores higher than 0.9 (IDP > 0.9) and less than 0.5 (IDP < 0.5). **c**. Domain enrichment analysis of intranuclear proteins with IDP scores higher than 0.9 by Pfam and Uniprot sequence feature. **d**. GO enrichment analysis of intranuclear proteins with IDP scores higher than 0.9.

To evaluate the subcellular localization of phase-separated proteins, we performed cellular component enrichment analysis on the 1467 intranuclear proteins. The result of enrichment revealed. As shown in Fig. 4d, these intranuclear proteins were highly enriched in proteins located in nucleolus, PML body and Cajal body. We also found that the molecular function terms RNA binding, DNA binding and protein binding were highly enriched in 1467 intranuclear proteins.

Proteins which are critical for forming phase separation belong to two archetypes ^1, 4, 6, 13^. The first type is characterized by multiple modular domains that interact with short linear motifs of other proteins, as well as RNAs and DNAs. The second type is characterized by the presence of IDRs that provide multiple weak adhesive sequence elements. The results of our sequential and functional analysis on 1467 intranuclear proteins indicate that candidates selected by IDeepPhase possess the typical characteristics of phase-separated proteins.

Cellular component enrichment analyses of 876 extranuclear proteins revealed that these proteins were significantly enriched to locate in P-bodies (Supplementary Fig. 4). However, the proteins were also significant to located in endosomes and lysosome (Supplementary Fig. 4), which indicates that proteins located in endosomes/lysosomes cannot be easily distinguished from components of membraneless organelles.

### Protein kinases potentially involved in phase separation

Post-translational modifications, particularly phosphorylation, are known to play crucial roles during condensation/dissolution of phase-separated compartments. For example, after being phosphorylated by kinase ZAP70, transmembrane protein LAT recruits several ligands such as protein SOS1, and subsequently forms the LAT complex that coalesces into T cell microclusters, enriching kinases while excluding phosphatases, subsequently activating downstream signaling pathways ^14^. Another example is the kinase DYRK3, which can regulate the mTORC pathway by regulating the condensation/dissolution of stress granules through its phosphorylation behavior ^18^. As shown in Fig. 5a, we found that 2343 proteins with IDP scores higher than 0.9 were preferentially phosphorylated (Fig. 5a). Especially, SOS1 and mTOR in the above two examples achieved IDP scores 0.998 and 0.956 respectively.

**Fig. 5.**
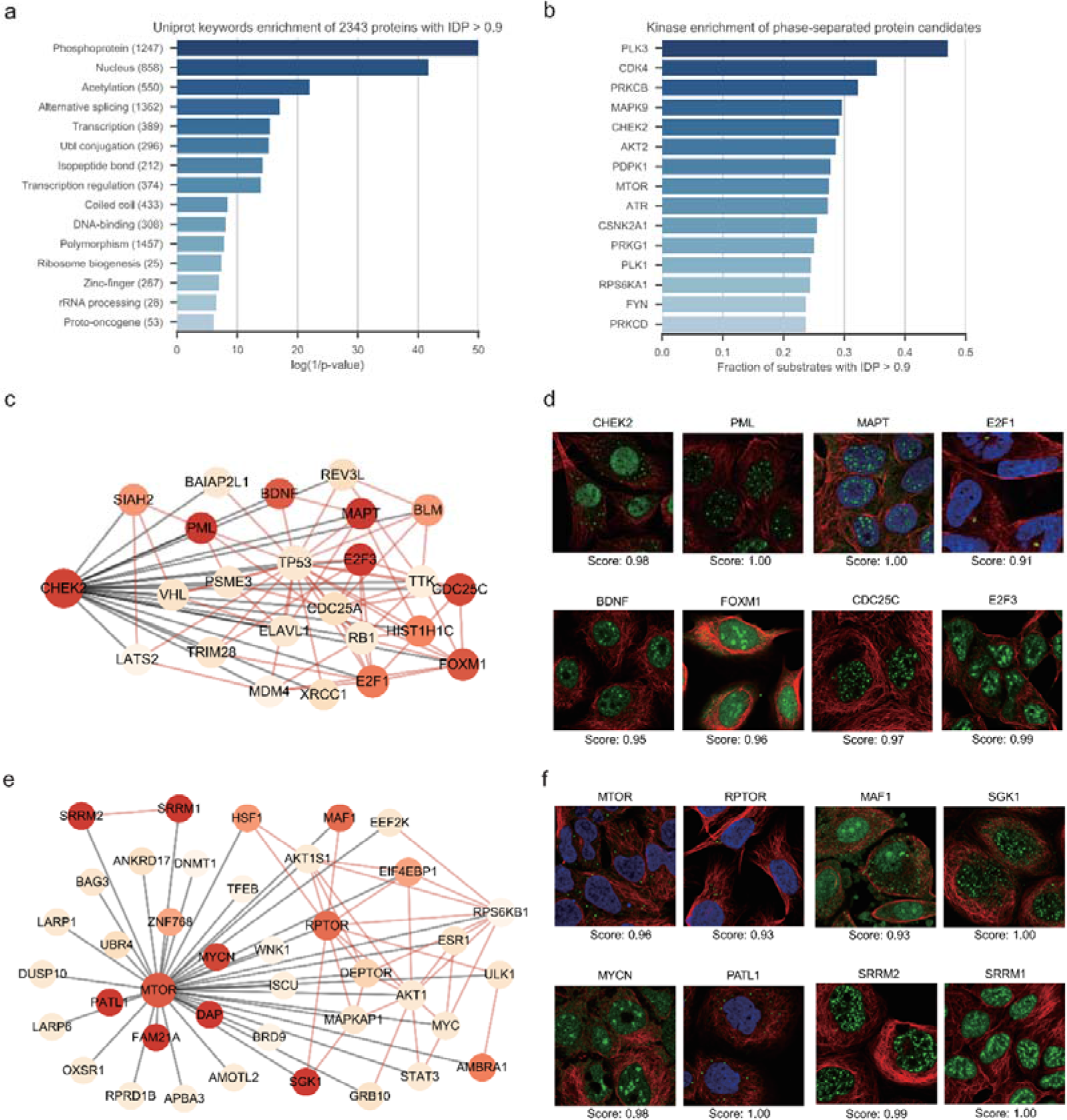
Preference of phosphorylation in proteins with IDP scores higher than 0.9. **a**. Enriched Uniprot keywords of proteins with IDP score higher than 0.9. **b**. For kinases with more than 15 scored substrates, the proportion of substrates with IDP scores higher than 0.9 were ranked. **c**.**d**. Substrate networks of CHEK2 (**c**) and mTOR (**d**). The node colors are correlated with IDP scores. Edges between kinases and substrates are denoted in grey, and edges within substrates are denoted in red. For each kinase, immunofluorescence images of eight substrates are shown.

For the 2343 proteins, 94% (2197/2343) possess phosphorylation sites, and 41% (897/2197) possess phosphorylation sites of known kinases. For kinases with more than 15 scored substrates, we calculated their proportion of substrates with IDP scores higher than 0.90 (Supplementary Table 5). As shown in Fig. 5b, the top 5 kinases were identified as PLK3, JNK2, PKCB, CDK4 and Chk2. One of these kinases, Chk2, is known to phosphorylate PML ^18,30,31^ and tau ^32,33,34^ (Fig. 5c). In addition to PML and tau, the other two substrates of Chk2, CDC25C and BDNF, achieved IDP scores of 0.97 and 0.95 (Fig. 5c) respectively, implying their potential to undergo polymerization. We also found that a considerable number of substrates of the kinase mTOR were involved in phase separation (Fig. 5d). mTOR is known to regulate phase separation of PGL granules during C. elegans embryogenesis through phosphorylation of PGL-1 and PGL-3 ^17^. We found that in human cells, mTOR and its substrates PRTOR, MAF1, SGK1, MYCN, PATL1, SRRM2 and SRRM1 formed droplets in IF images (Fig. 5d), suggesting that phase separation dose crucial to the mTOR pathway.

## Discussion

In this study, we established a methods called IDeepPhase which discriminated IF images with droplet structures. Using IDeepPhase on the proteome-scale IF images, a list of proteins which have displayed as droplet structures were obtained. Different from sequence analysis methods, candidate phase-separated proteins selected by IDeepPhase have exhibited multimerization potential. The formation of droplet structures depended on different conditions ^1, 6^. As protein SOS1 in Fig. 2d and protein EWSR1 in Supplementary Fig. 2 displayed, they only displayed droplet structures in a part of IF images. The antibody and cell line information of each IF image were included in Supplementary Table 3 to aid in the selection of antibody and cell line for subsequent phase separation study.

Besides phase-separated proteins, proteins located in extranuclear membrane-bounded organelles like endosomes or lysosomes display spherical structures on IF images as well. Though the size of endosomes or lysosomes are slightly different from membraneless organelles; endosomes and lysosomes vary in size from tens of nanometers to hundreds of nanometers, while membraneless organelles are in micron scale ^1^. Proteins located in endosomes and lysosome still hard to be discriminated from those proteins located in membraneless organelle components. Proteins which were assigned to endosomes and lysosomes by Swiss-Prot database were labeled in Supplementary Table 3. While for 1467 proteins located in the nucleus where no classic membrane-bound organelles have been reported, their sequential and functional preferences are in agreement with features of phase-separated proteins, which indicated that intranuclear candidates are more reliable.

The importance of phosphorylation in phase separation is well established. It is interesting to answer the question that those potential components of phase-separated compartments prefer to be regulated by which kinases. A list of kinases whose substrates displayed preference of high IDP scores were obtained. Take kinase Chk2 in this list as an example, multiple evidences have proved that Chk2 kinase phosphorylate PML on Ser-117 and colocalize in PML nuclear bodies ^35,36,37,38,39^. Furthermore, it has been found that Chk2’s interaction with proteasome activator REGγ is necessary for REGγ’s regulative effect on the number of PML nuclear bodies ^36^. However, the functions of Chk2 in phase-separated compartments have not been clear displayed. The kinases uncovered in this study provide new ways of understanding well-established phosphorylation signaling pathways.

## Methods

### Collection of proteins involved in phase separation

3569 articles were extracted from PubMed on Oct 21, 2018, using the keyword ((((((“phase transition”[Title/Abstract]) OR “phase separation”[Title/Abstract]) OR membraneless organelles[Title/Abstract]) OR biomolecular condensate[Title/Abstract]) AND protein) AND cell), and these articles were manually reviewed. As a result, 96 human proteins were confirmed to be involved in phase-separation. (Supplementary Table 1).

Cell Atlas of Human Protein Atlas database (https://www.proteinatlas.org/humanproteome/cell) ^21^ provides antibody-based profiling by immunofluorescence confocal microscopy for 12073 proteins with available antibodies. In total, 83673 IF images by different antibodies in different cell lines were downloaded and stored for subsequent analysis.

90 of 96 collected human phase-separated proteins have available IF images, and they were mapped to 812 IF images. The formation of phase separation is conditional dependent, and those experimentally verified phase-separated proteins cannot display as spherical droplets in all IF images. Therefore, 274 of 812 images which display as droplets with higher fluorescence intensity than backgrounds were selected for further analysis (Supplementary Table 2).

### Convolutional neural network

As described in Supplementary Fig. 1, Our CNN model contained seven layers with trainable weights; the first five were convolutional and the remaining two were fully-connected. Convolutional layers were activated with RELU function and connected with max-pooling layers which pooling each 2*2 pixels into 1 pixel; the last max-pooling layer was linked with a global max-pooling layer, which pool a 14*14*64 tensor into a vector of length 64. Then the vector was connected with a fully connected layer with 64 neurons and RELU activation, a dropout layer with dropout rate equal to 0.2, and a fully connected layer with one neuron and sigmoid activation. The output of the sigmoid function was a value range from 0 to 1, which represents the possibility of image classification. We built our model with Keras 2.2 and TensorFlow 1.12.

### Two-stage training process of IDeepPhase

Two-stage training was applied in the training process (Fig. 1a). In the first stage, the CNN network was trained with the positive training set which includes 195 images and an equal size negative training set which were randomly selected from the 83673 IF images after filtering out positive samples. The remaining 79 IF images and an equal size images in negative set were used as test dataset. To augment the size of the training and test datasets, each image (3, 2048, 2048) was cropped into 4 non-overlapping pieces (3, 512, 512), which increased the number of IF images by 4 times. The sample sizes of training set and test set are 1560 (780 positive VS 780 negative samples) and 632 (316 positive VS 316 negative samples) respectively. ImageDataGenerator module from Keras library was applied to generate rotated and flipped images. The CNN model was training with the cropped training sets. Adam optimizer in Keras library were applied to train the CNN with default parameters and batch size equal to 64 for 50 epochs. The obtained CNN classifier was use to score the cropped IF image in the set. For each original IF image in the test set, the highest score of four cropped images was treated as the final score. The AUC of the obtained CNN classifier on the test dataset was 0.72.

In the second stage, each download IF image was cropped into 4 non-overlapping pieces and finally 334692 cropped images were obtained. We applied the CNN classifier established in the first stage on all the 334692 cropped images, and top 1000 scored images non-overlapped with the positive and negative sample sets were manually checked and labeled. 934 manually labeled cropped images (461 positive samples and 473 negative samples) were added to the positive and negative training set of the first stage. Then the CNN network was retrained with this extended training set, and the test dataset was the same as the first stage. The AUC on the test set was 0.90 in the second stage. We further set signals from red (microtubules) and blue (nucleus) channels to zero, and using only green (antibody) channel; the AUC on the same test set was 0.91. The final model used in IDeepPhase was the green-only model.

### Location, function and phosphorylation sites annotations

Subcellular localization of each image was extracted from Cell Atlas of the Human Protein Atlas database. Proteins with location annotations include nucleoli, nucleoli fibrillar center, nuclear bodies, nuclear speckles, and nucleus were denoted as intranuclear proteins (Fig. 4a). Otherwise, the proteins were denoted as extranuclear proteins (Fig. 4a).

Enriched GO terms, Pfam domains, and Uniprot keywords were identified by DAVID 6.8 ^40^.

The phosphorylation site dataset and kinase–substrate interaction dataset was downloaded from the PhosphoSitePlus database, which included 237523 phosphorylation sites and 10266 kinase–substrate relations ^41^.

### Code availability

Source code are available at https://github.com/cheneyyu/IDeepPhase.

## Acknowledgments

We thank Dr. Jun Qin (National Center for Protein Sciences, China) for helpful suggestions and Dr. Torsten Juelich (Peking University, China) for linguistic assistance during preparation of our manuscript. This work was supported by the National Key Research and Development Program of China [2018YFA0507504]; the National Natural Science Foundation of China [61773025].

## Author Contributions

C.Y. and T.L. conceived the study; C.Y., and T.L. designed the research; C.Y. and B.S. performed the research; B.S., Q.H., and K.Y. collected and labeled the data; C.Y., B.S. and M.S. analyzed data; Y.C. and C.W. contributed to the data analysis; and C.Y., B.S. and T.L. wrote the paper with input from all other authors.

## Competing interests

The authors declare no competing interests.

## Supplementary Files

Supplementary Fig. 1 Convolutional Neural Network architecture of IDeepPhase.

Supplementary Fig. 2 IDeepPhase scores and immunofluorescence images of EWSR1 in 3 cell lines: A-431, U-2 OS and U-251MG.

Supplementary Fig. 3 IDeepPhase scores and immunofluorescence images of five PML body components, ATRX, MRE11, HIPK3, RPAIN and TDP2.

Supplementary Fig. 4 GO enrichment analysis of extranuclear proteins with IDP scores higher than 0.9.

Supplementary Table 1 Summary of collected known phase-separated proteins.

Supplementary Table 2 Summary of images used in two stages of training and validation.

Supplementary Table 3 Summary of IDeepPhase scores in 12073 proteins with information of related cell lines, antibodies, and locations.

Supplementary Table. 4 Summary of known phase-separated proteins in different bodies.

Supplementary Table 5 Summary of IDeepPhase scores in kinase substrate interactions.

Supplementary Table 6 Summary of IDeepPhase scores in 83673 images with information of related cell lines, antibodies, and locations.

## References

1. Banani SF, Lee HO, Hyman AA, Rosen MK. Biomolecular condensates: organizers of cellular biochemistry. Nat Rev Mol Cell Biol 18, 285–298 (2017).

2. Hyman AA, Weber CA, Julicher F. Liquid-liquid phase separation in biology. Annu Rev Cell Dev Biol 30, 39–58 (2014).

3. Brangwynne CP, et al. Germline P Granules Are Liquid Droplets That Localize by Controlled Dissolution/Condensation. SCIENCE 324, 1729–1732 (2009).

4. Boeynaems S, et al. Protein Phase Separation: A New Phase in Cell Biology. Trends in Cell Biology 28, 420–435 (2018).

5. Boulay G, et al. Cancer-Specific Retargeting of BAF Complexes by a Prion-like Domain. Cell 171, 163–178 e119 (2017).

6. Shin Y, Brangwynne CP. Liquid phase condensation in cell physiology and disease. Science 357, (2017).

7. Harmon TS, Holehouse AS, Rosen MK, Pappu RV. Intrinsically disordered linkers determine the interplay between phase separation and gelation in multivalent proteins. Elife 6, (2017).

8. Oates ME, et al. D2P2: database of disordered protein predictions. Nucleic Acids Research 41, D508–D516 (2012).

9. Potenza E, Domenico TD, Walsh I, Tosatto SCE. MobiDB 2.0: an improved database of intrinsically disordered and mobile proteins. Nucleic Acids Research 43, D315–D320 (2015).

10. Vernon RM, et al. Pi-Pi contacts are an overlooked protein feature relevant to phase separation. Elife 7, (2018).

11. Lancaster AK, Nutter-Upham A, Lindquist S, King OD. PLAAC: a web and command-line application to identify proteins with prion-like amino acid composition. Bioinformatics 30, 2501–2502 (2014).

12. Stanislawski J, Kotulska M, Unold O. Machine learning methods can replace 3D profile method in classification of amyloidogenic hexapeptides. BMC Bioinformatics 14, 21 (2013).

13. Alberti S, Gladfelter A, Mittag T. Considerations and Challenges in Studying Liquid-Liquid Phase Separation and Biomolecular Condensates. Cell 176, 419–434 (2019).

14. Su X, et al. Phase separation of signaling molecules promotes T cell receptor signal transduction. SCIENCE 352, 595–599 (2016).

15. Aulas A, et al. Stress-specific differences in assembly and composition of stress granules and related foci. J Cell Sci 130, 927–937 (2017).

16. Bernardi R, et al. PML inhibits HIF-1alpha translation and neoangiogenesis through repression of mTOR. Nature 442, 779–785 (2006).

17. Zhang G, Wang Z, Du Z, Zhang H. mTOR Regulates Phase Separation of PGL Granules to Modulate Their Autophagic Degradation. Cell 174, 1492–1506 e1422 (2018).

18. Berchtold D, Battich N, Pelkmans L. A Systems-Level Study Reveals Regulators of Membrane-less Organelles in Human Cells. Mol Cell 72, 1035–1049 e1035 (2018).

19. Alberti S, Saha S, Woodruff JB, Franzmann TM, Wang J, Hyman AA. A User’s Guide for Phase Separation Assays with Purified Proteins. J Mol Biol 430, 4806–4820 (2018).

20. Sullivan DP, et al. Deep learning is combined with massive-scale citizen science to improve large-scale image classification. Nature Biotechnology 36, 820–828 (2018).

21. Thul PJ, et al. A subcellular map of the human proteome. Science 356, (2017).

22. Protter DSW, et al. Intrinsically Disordered Regions Can Contribute Promiscuous Interactions to RNP Granule Assembly. Cell Rep 22, 1401–1412 (2018).

23. Hubstenberger A, et al. P-Body Purification Reveals the Condensation of Repressed mRNA Regulons. Molecular Cell 68, 144–157.e145 (2017).

24. Feric M, et al. Coexisting Liquid Phases Underlie Nucleolar Subcompartments. Cell 165, 1686–1697 (2016).

25. Fong KW, et al. Whole-genome screening identifies proteins localized to distinct nuclear bodies. J Cell Biol 203, 149–164 (2013).

26. Youn JY, et al. High-Density Proximity Mapping Reveals the Subcellular Organization of mRNA-Associated Granules and Bodies. Mol Cell 69, 517–532 e511 (2018).

27. Jain S, Wheeler JR, Walters RW, Agrawal A, Barsic A, Parker R. ATPase-Modulated Stress Granules Contain a Diverse Proteome and Substructure. Cell 164, 487–498 (2016).

28. Nover L, Scharf KD, Neumann D. Cytoplasmic heat shock granules are formed from precursor particles and are associated with a specific set of mRNAs. Molecular and cellular biology 9, 1298–1308 (1998).

29. Kirmitzoglou I, Promponas VJ. LCR-eXXXplorer: a web platform to search, visualize and share data for low complexity regions in protein sequences. Bioinformatics 31, 2208–2210 (2015).

30. Bernardi R, Pandolfi PP. Structure, dynamics and functions of promyelocytic leukaemia nuclear bodies. Nat Rev Mol Cell Biol 8, 1006–1016 (2007).

31. Dellaire G, Eskiw CH, Dehghani H, Ching RW, Bazett-Jones DP. Mitotic accumulations of PML protein contribute to the re-establishment of PML nuclear bodies in G1. J Cell Sci 119, 1034–1042 (2006).

32. Patel A, et al. A Liquid-to-Solid Phase Transition of the ALS Protein FUS Accelerated by Disease Mutation. Cell 162, 1066–1077 (2015).

33. Ambadipudi S, Biernat J, Riedel D, Mandelkow E, Zweckstetter M. Liquid-liquid phase separation of the microtubule-binding repeats of the Alzheimer-related protein Tau. Nat Commun 8, 275 (2017).

34. Wegmann S, et al. Tau protein liquid-liquid phase separation can initiate tau aggregation. EMBO J 37, (2018).

35. Stolz A, Ertych N, Bastians H. Tumor suppressor CHK2: regulator of DNA damage response and mediator of chromosomal stability. Clin Cancer Res 17, 401–405 (2011).

36. Zannini L, Buscemi G, Fontanella E, Lisanti S, Delia D. REGgamma/PA28gamma proteasome activator interacts with PML and Chk2 and affects PML nuclear bodies number. Cell Cycle 8, 2399–2407 (2009).

37. di Masi A, et al. PML nuclear body disruption impairs DNA double-strand break sensing and repair in APL. Cell Death Dis 7, e2308 (2016).

38. Yang S, Kuo C, Bisi JE, Kim MK. PML-dependent apoptosis after DNA damage is regulated by the checkpoint kinase hCds1/Chk2. Nat Cell Biol 4, 865–870 (2002).

39. Varadaraj A, et al. Evidence for the receipt of DNA damage stimuli by PML nuclear domains. J Pathol 211, 471–480 (2007).

40. Huang DW, Sherman BT, Lempicki RA. Systematic and integrative analysis of large gene lists using DAVID bioinformatics resources. Nature Protocols 4, 44 (2008).

41. Hornbeck PV, Zhang B, Murray B, Kornhauser JM, Latham V, Skrzypek E. PhosphoSitePlus, 2014: mutations, PTMs and recalibrations. Nucleic Acids Res 43, D512–520 (2015).

